# Effects of canagliflozin and irbesartan on renal fibrosis in Dahl salt-sensitive rats

**DOI:** 10.1101/2022.12.27.522015

**Authors:** Jianlong Zhai, Zhongli Wang, Tingting Zhang, Lili He, Sai Ma, Qingjuan Zuo, Guorui Zhang, Xinyu Wang, Yifang Guo

## Abstract

Hypertension is one of the major contributors to cardiovascular and chronic kidney disease (CKD). Sodium-glucose cotransporter 2 (SGLT2) inhibitors and angiotensin receptor blockers (ARBs) have become the preferred treatment for patients with CKD. However, the renoprotective effects of the combined therapy of the two drugs on hypertensive renal fibrosis are still largely understood. The aim of this study was to compare the antifibrotic effects of canagliflozin, with or without irbesartan, in the kidneys of Dahl salt-sensitive (Dahl SS) rats on a high salt (HS) diet. After the preconditioning stage, Dahl SS rats (n = 47) were divided into 5 experimental groups as follows: low salt control (n=7), HS control (n=10), high salt with canagliflozin (n=10), high salt with irbesartan (n=10), and high salt with canagliflozin plus irbesartan (n=10). Mean food and water intake, body weight (BW), and systolic blood pressure (SBP) were measured during the whole experimental period. After 12 weeks, the rats were euthanized, and the kidneys were excised for histomorphometric evaluation and immunohistochemical evaluation. An HS diet increased SBP, renal fibrosis, expression of fibrotic protein factors, and TGF-β/Smad2/3 pathway compared to the LS group. We found that irbesartan reduced SBP and slowed the loss of renal function. Canagliflozin significantly reduced BW and renal fibrosis and downregulated the TGF-β/Smad2/3 pathway. The combined therapy showed better renoprotection in all outcome parameters. In conclusion, these results indicate that canagliflozin and irbesartan exert different benefits on nephroprotection in salt-sensitive hypertensive rats.

## Introduction

Among the host of factors leading to essential hypertension, high dietary salt intake, always leading to salt-sensitive hypertension (SSHT), is a key environmental factor [1]. Although restricted salt intake is beneficial, people continue to consume excessive salt in some areas, especially in North China [2]. Hypertension with chronic kidney disease (CKD) is caused by various factors, among which salt sensitivity and increased activity of the renin-angiotensin-aldosterone and sympathetic nervous systems might be most important [3]. Therefore, we pay close attention to the study of kidney damage in SSHT.

Salt-sensitive hypertensive individuals have a higher plasma volume, total peripheral resistance, and blood pressure, mainly through the retention of water and sodium by high salt intake [4]. Renal fibrosis is a characteristic kidney pathogenesis of SSHT [5]. Transforming growth factor-β (TGF-β), mainly regulated via Smad proteins, is the most important profibrotic cytokine in renal fibrosis [6]. Dahl salt-sensitive (Dahl SS) rats develop SSHT and chronic ischemic nephropathy with a high salt (HS) diet, which has been demonstrated to induce proteinuria, glomerulosclerosis, interstitial fibrosis, and eventually progressive loss of renal function [5].

Sodium-glucose cotransporter 2 (SGLT2) inhibitors are a new category of antidiabetic drugs [7] that inhibit glucose reabsorption in renal proximal tubules, leading to increased urinary glucose excretion and lower plasma glucose levels [8]. Multicenter clinical trials [9] demonstrated that SGLT2 inhibitors can improve renal function and reduce the risk of major kidney events in T2D patients. Moreover, SGLT2 inhibitors are promising as first-line therapies for nondiabetic CKD [10]. However, the effect of SGLT2 inhibitors on renal fibrosis of SSHT has not been well-studied. Because angiotensin II receptor blockers (ARBs) are the basic treatment for CKD, the aims of our study were to explore the effects of canagliflozin and irbesartan on renal injury in Dahl SS rats on an HS diet.

## Results

### Changes in physiological data

We first confirmed the differences in BW and the other physiological parameters among the five groups during the 12-week experiment (Fig. 1). After treatment for 4 weeks, BW was lower in the HS group than in the LS group, was lower in the HS + CANA group than in the HS + IRB group, and was lower in the HS + CANA + IRB group than in the HS+IRB group. After treatment for 4 weeks, mean food intake (MFI) was lower in the HS + CANA group than in the HS + IRB group and was lower in the HS + CANA+IRB group than in the HS + IRB group. After treatment for 6 weeks, the mean water intake (MWI) was higher in the HS group than in the LS group, higher in the HS + CANA group than in the HS + IRB group, and higher in the HS + CANA + IRB group than in the HS + IRB group. After treatment for 5 weeks, SBP was higher in the HS group than in the LS group, lower in the HS + IRB group than in the HS group, and lower in the HS + IRB + CANA group than in the HS group. At 13 weeks of age, FBG was lower in the HS + CANA group than in the LS group, lower in the HS + CANA group than in the HS group, lower in the HS + CANA + IRB group than in the LS group, and lower in the HS + CANA + IRB group than in the HS group (Fig. 2A). No significant differences in RBG were found among the groups (Fig. 2B).

**Fig. 1.**
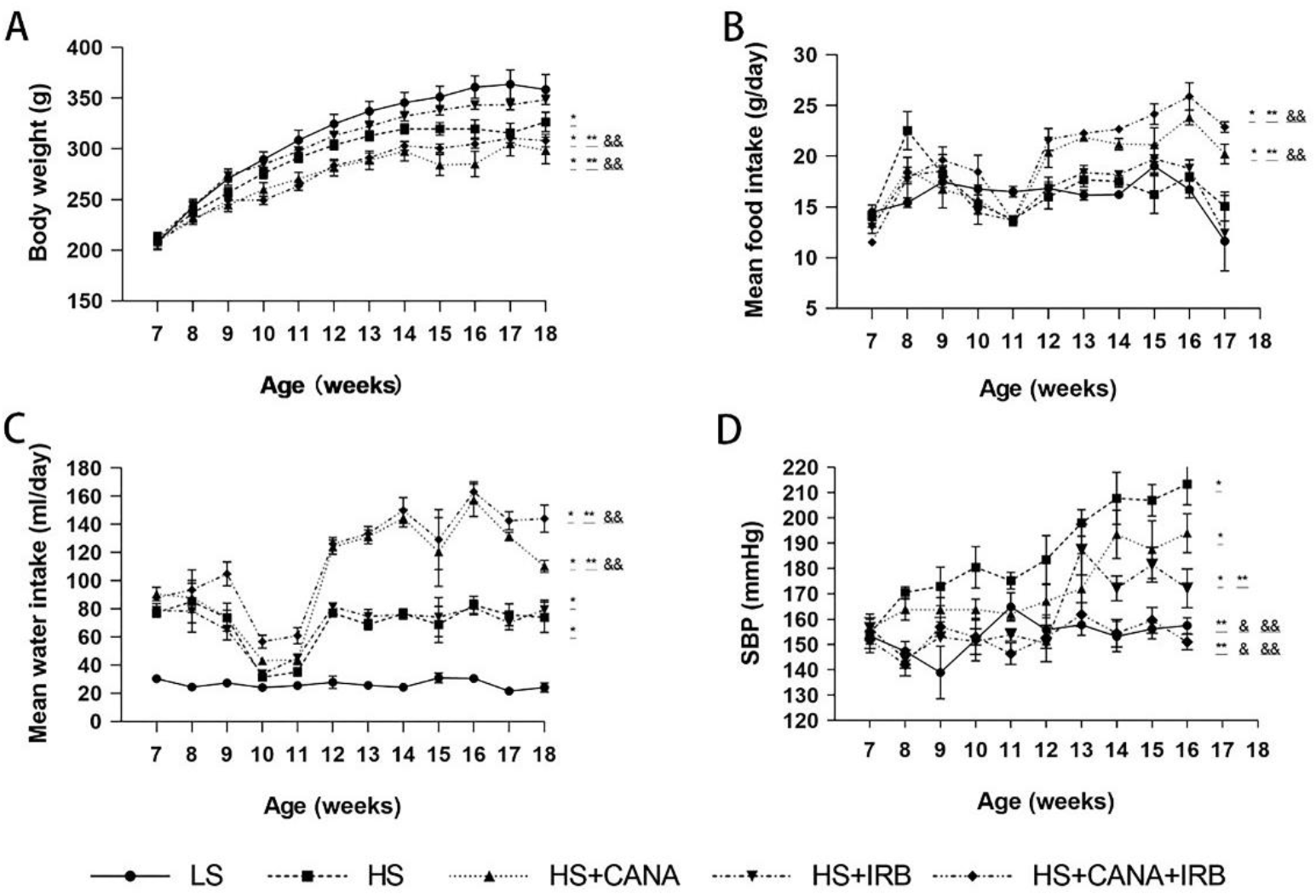
Time courses of body weight, mean food and water intake, and SBP for rats of the five experimental groups. A–D Changes in body weight, mean food intake, mean water intake, and SBP, respectively. All data are means ± SEM (n = 7, 11, 9, 9, and 9 rats in A–D for LS, HS, HS+CANA, HS+IRB and HS+CANA+IRB groups, respectively). *P < 0.05 vs. LS, **P < 0.05 vs. HS, &P < 0.05 vs. HS+CANA, &&P < 0.05 vs. HS+IRB.

**Fig. 2.**
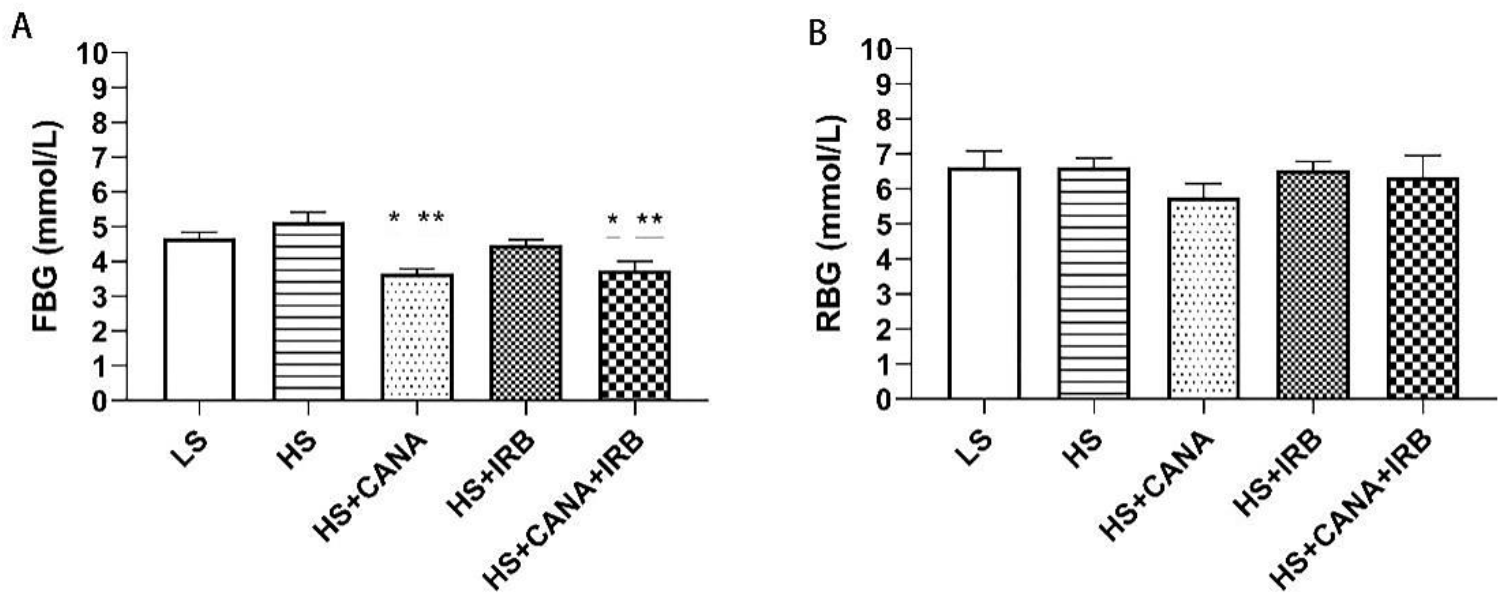
Fasting blood glucose and random blood glucose for rats of the five experimental groups at 13 weeks of age. *P < 0.05 vs. LS, **P < 0.05 vs. HS.

### Physiological and biochemical data at the end of the experiment

At the end of the experiment, we examined the effects of different treatments on physiological and biochemical data among the five groups (Table 1). Compared with the LS group, the HS group, the HS+CANA group, and the HS+CANA+IRB group had lower BW. Compared with the HS+IRB group, the HS group and the HS+CANA group had lower BW. BW was lower in the HS+CANA+IRB group than in the HS+IRB group. There were no significant differences in FBG among the five groups. Compared with the HS group, the LS group, the HS+IRB group, and the HS+CANA+IRB group had lower SBP. SBP in the HS+CANA+IRB group was lower than that in the HS+CANA group. Compared with the LS group, the HS group, the HS+IRB group, and the HS+CANA group had a lower MFI. The MFI in the HS+ CANA+ IRB group was higher than that in the HS group, the HS+CANA group, and the HS+IRB group. Compared with the LS group, the other four groups had a higher MWI. The heart rate (HR) in the HS+CANA+IRB group was lower than that in the HS group. Compared with the LS group and the HS group, the urine volume of the HS+CANA group significantly increased, while the level of urine volume was decreased by irbesartan in the HS+CANA+IRB group.

**Table 1.** Physiological and biochemical parameters of the rats at the end of the 12-week experiment.

| Parameter | LS | HS | HS+CA | HS+IRB | HS+CA+IRB |
| --- | --- | --- | --- | --- | --- |
| BW (g) | 363.5 ± 35.8 | 316.1 ± 38.1* | 297.9 ± 43.1* | 346.6 ± 17.8**,# | 306.9 ± 16.8*,## |
| SBP (mmHg) | 175.8 ± 28.3 | 213.4 ± 18.4* | 194.0 ± 19.0 | 172.2 ± 18.7** | 151.0 ± 7.0**,# |
| LKW (g) | 1.31 ± 0.09 | 1.48 ± 0.15 | 1.33 ± 0.45 | 1.56 ± 0.04 | 1.42 ± 0.14 |
| RKW (g) | 1.29 ± 0.04 | 1.45 ± 0.21 | 1.44 ± 0.27 | 1.42 ± 0.05 | 1.276 ± 0.004 |
| MFI (g/day) | 22.7 ± 2.4 | 15.6 ± 0.7* | 18.2 ± 5.2* | 18.8 ± 0.8 | 22.8 ± 0.4**,#,## |
| MWI (ml/day) | 24.1 ± 7.4* | 73.90 ± 24.0* | 91.2 ± 43.2* | 79.3 ± 15.3* | 123.7 ± 48.7*,**,## |
| HR (bpm) | 389.2 ± 43.1 | 426.2 ± 54.3 | 381.7 ± 31.0 | 385.0 ± 27.6 | 375.2 ± 34.9** |
| FBG (mmol/L) | 11.9 ± 1.0 | 11.1 ± 0.2 | 14.5 ± 1.4* | 14.1 ± 1.1 | 14.5 ± 1.2* |
| UV (ml) | 19.5 ± 8.5 | 51.0 ± 7.8 | 137.0 ± 21.8*,** | 49.3 ± 10.3# | 91.0 ± 33.2* |
| 24-h UTP (mg) | 23.2 ± 12.7 | 82.6 ± 20.8* | 50.9 ± 9.3 | 37.7 ± 14.9** | 19.2 ± 3.0** |
| FENa (%) | 0.17 ± 0.05 | 2.26 ± 0.46* | 2.73 ± 0.94* | 0.31 ± 0.24**,# | 1.00 ± 1.04**,# |
| Scr (mmol/L) | 43.4 ± 2.5 | 51.7 ± 8.7 | 42.2 ± 11.6 | 37.9 ± 3.7** | 31.1 ± 7.3*,**,# |
| BUN (mmol/L) | 5.9 ± 1.1 | 8.1 ± 1.4 | 9.0 ± 1.2 | 7.1 ± 0.8 | 11.1 ± 0.9 |
| Cys-C (mg/L) | 3.74 ± 0.43 | 4.22 ± 0.12 | 4.45 ± 0.14 | 3.78 ± 0.30 | 4.45 ± 0.10 |
| KIM-1 (ng/ml) | 2.04 ± 0.10 | 1.83 ± 0.08 | 2.08 ± 0.07 | 1.85 ± 0.10 | 1.90 ± 0.06 |
| UG (mmol/L) | 1.6 ± 0.5 | 1.0 ± 0.1 | 185.9 ± 21.0*,** | 10.8 ± 8.0# | 413.9 ± 49.3*,**,# |
| UNa (mmol/L) | 25.2 ± 3.2 | 214.4 ± 9.4* | 176.2 ± 6.3* | 85.1 ± 38.1*,**,# | 112.6 ± 17.0*,**,# |
| UCr (μmol/L) | 5731.5 ± 380.7 | 3811.0 ± 349.0 | 1420.3 ± 204.7*,** | 4747.6 ± 959.7# | 3751.3 ± 521.5*,# |
| UUN (mmol/L) | 181.1 ± 9.3 | 123.0 ± 9.0* | 96.8 ± 9.4* | 155.6 ± 18.5# | 113.6 ± 12.9*,## |

Data are means ± SEMs (n = 7, 9, 8, 9, and 10 rats for physiological data and n = 5, 5, 5, 5, and 5 rats for biochemical data of LS, HS, HS+CA, HS+IRB and HS+CA+IRB groups, respectively). \*P < 0.05 vs. LS. \*\*P < 0.05 vs. HS. #P < 0.05 vs. HS+CA. ##P < 0.05 vs. HS+IRB. BW: Body weight; FBG: Fasting blood glucose; SBP: Systolic blood pressure; LKW: Left kidney weight; RKW: Right kidney weight; MFI: Mean food intake; MWI: Mean water intake; HR: Heart rate; 24-h UTP: 24-hour urinary total protein; UV: Urine volume; FENa: Fractional excretion of urinary sodium; Scr: Serum creatinine; BUN: Blood urea nitrogen; Cys-C: Cystatin C; KIM-1: Kidney injury molecule 1; UG: Urine glucose; UNa: Urine sodium; UCr: Urine creatinine; UUN: Urine urea nitrogen.

To assess markers of renal injury in the rats, we next measured several important indexes of kidney function biomarkers (Table 1). Compared with the LS group, the 24 h urinary protein of the HS group significantly increased. After treatments, there was a significant decrease in the 24 h urinary protein in the HS+IRB group and the HS+CANA+IRB group. The 24 h urinary protein of the HS+CANA group also decreased, but the difference was not statistically significant. The same was observed in serum creatinine (Scr) and fractional excretion of urinary sodium (FENa). There were no significant changes in urea nitrogen, cystatin C, or kidney injury molecule 1 in the five different groups.

### Renal histological changes

We then performed a series of histochemical staining, including H&E, Masson, and TGF-β1 staining, to evaluate the pathological changes in the rat kidneys of the five different groups. The H&E staining results demonstrate that compared to the LS group, morphological abnormalities of glomeruli, dilation of tubules, and epithelial cell edema were observed in the HS group, which were markedly attenuated in the HS+CANA, HS+IRB, and HS+CANA+IRB groups (Fig. 3A). The improvement of morphologic lesions was more obvious in the HS+CANA+IRB group. Masson staining (Fig. 3B) and TGF-β1 IF staining (Fig. 3C) were analyzed by Image 8.0 software. As shown by Masson staining (Fig. 3D), the interstitial fibrosis of the HS group was more evident than that of the LS group. The collagen deposition in the HS+CANA, HS+IRB and HS+CANA+IRB groups was markedly alleviated. The combined treatment group had a more beneficial effect. We used IF to analyze the expression levels of TGF-β1 in the kidneys of rats (Fig. 3C). The results showed that the expression changes in TGF-β1 were similar to the changes in collagen deposition in rat kidney tissue in the five different groups (Fig. 3E). Taken together, these results indicate that canagliflozin, irbesartan, and the combined treatment may improve renal fibrosis caused by an HS diet in Dahl SS rats, possibly through the TGF-β1/Smad2/3 pathway.

**Fig. 3.**
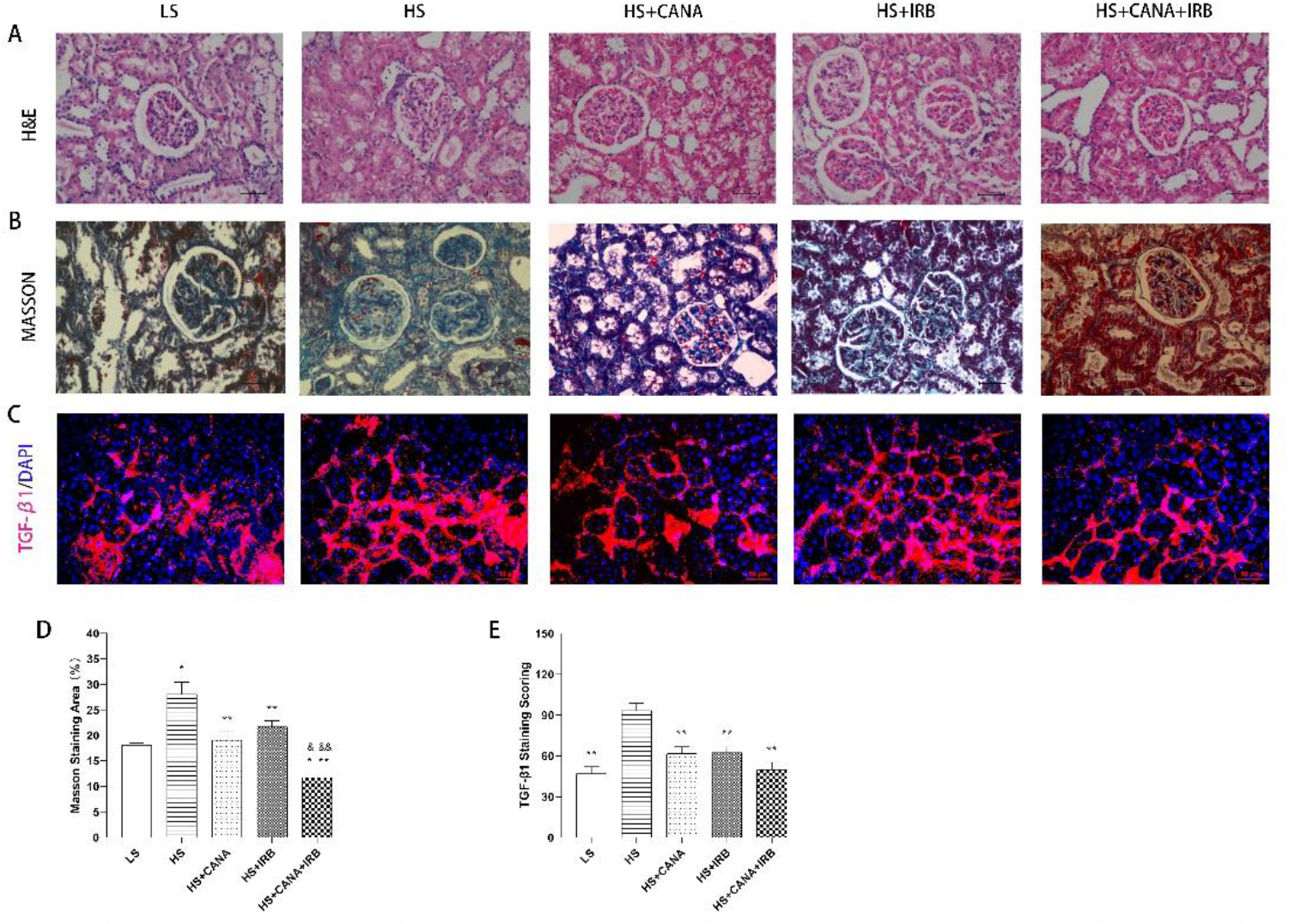
Renal histological damage in the different groups. (A) Hematoxylin and Eosin staining (Original magnification ×400). (B) Masson staining (Original magnification ×400). (C) Representative immunofluorescence images showing the induction of TGF-β1 protein in the renal cortex (Original magnification ×400). Slides are counterstained with DAPI for nuclear visualization. (D) The percentage of Masson-positive area (blue) relative to the whole area of Masson staining specimens in the different groups (one-way ANOVA, *p < 0.05 vs LS, **p < 0.05 vs HS, &p < 0.05 vs HS+CANA, &&p < 0.05 vs HS+IRB). (E) The percentage of TGF-β1-positive area (red) relative to the whole area of immunofluorescence staining specimens in the different groups (one-way ANOVA, *p < 0.05 vs LS, **p < 0.05 vs HS, &p < 0.05 vs HS+CANA, &&p < 0.05 vs HS+IRB). The values are expressed as mean ± SEM.

### Expression of fibrotic protein factors and the TGF-β/Smad2/3 pathway

To further confirm the effect of canagliflozin and irbesartan on renal fibrosis and the TGF-β1/Smad2/3 pathway, a WB assay was performed. Renal fibrosis is characterized by the production of collagen and the activation of α-SMA. Subsequently, the expression of collagen I, and α-SMA was determined by WB. The results showed that an HS diet significantly increased the protein expression of collagen I (Fig. 4B), and α-SMA (Fig. 4C) in the kidneys of Dahl SS rats, whereas these levels were downregulated by the HS+CANA group and the HS+CANA+IRB group. However, these fibrotic protein levels in the HS+IRB group were not significantly decreased. As shown in Figure 4, compared with the LS group, the HS group significantly upregulated the protein expression of TGF-β1 (Fig. 4D), Smad2/3 (Fig. 4E), and p-Smad2/3 (Fig. 4F), whereas these changes were reversed by canagliflozin and irbesartan. Similarly, additive effects were observed in the combined treatment group. In addition, compared with that in the LS group, the expression of Smad7 in the HS group (Fig. 4G) was significantly downregulated, whereas the expression of Smad7 in the HS+CANA, HS+IRB, and HS+CANA+IRB groups was dramatically upregulated. Taken together, these findings suggested that compared with irbesartan, canagliflozin may better alleviate renal fibrosis caused by an HS diet in Dahl SS rats, possibly through inhibiting the TGF-β1/Smad2/3 pathway.

**Fig. 4.**
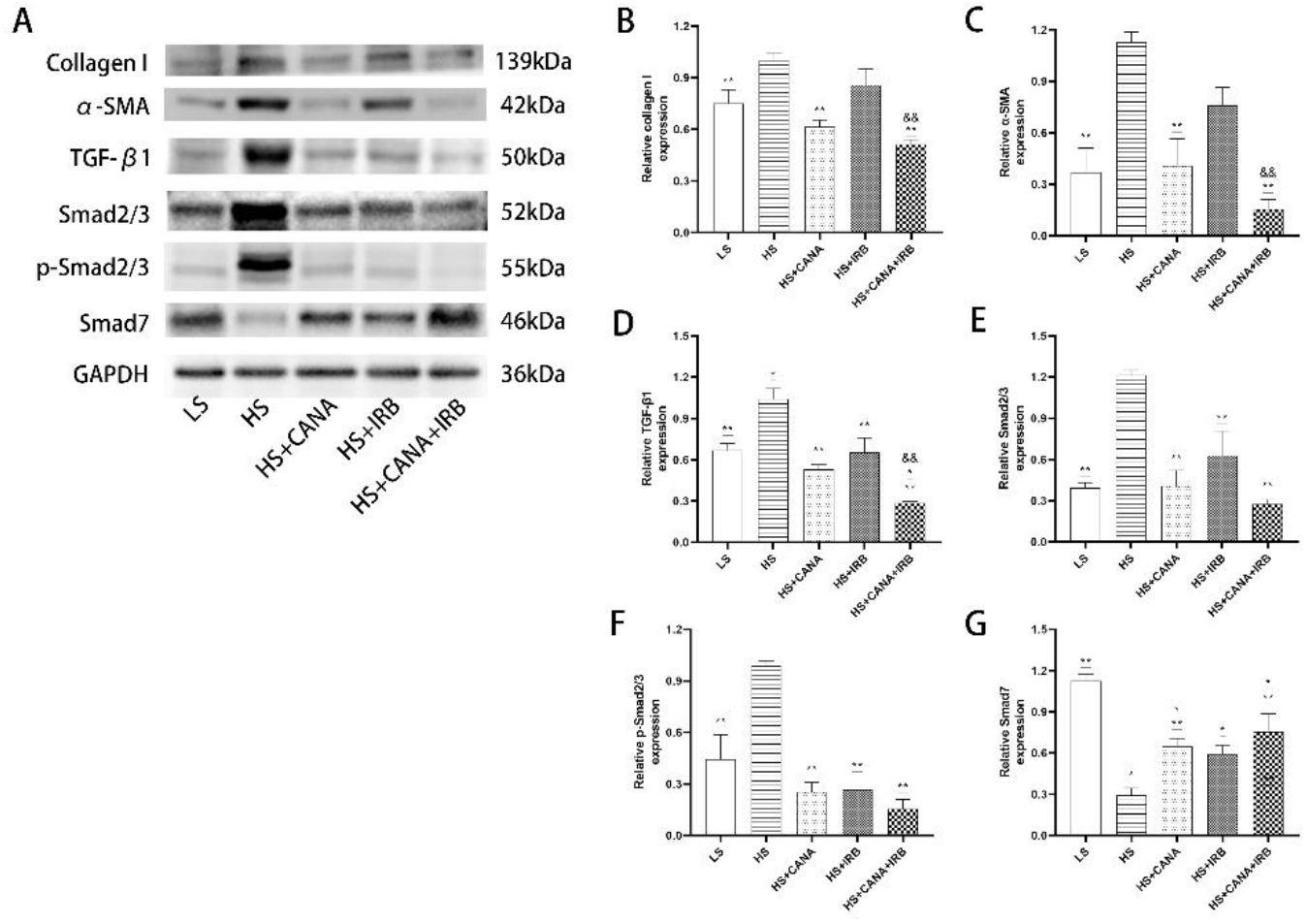
Effect on renal fibrotic factors and the TGF-β/Smad pathway in the five different groups. A The protein expressions of fibronectin, collagen I, α-smooth muscle actin (α-SMA), TGF-β1, Smad2/3, p-Smad2/3, Smad7 and glyceraldehyde-3-phosphate dehydrogenase (GAPDH) in the five different groups were measured by western blot. The relative expression levels of collagen I (B), α-SMA (C), TGF-β1 (D), Smad2/3 (E), p-Smad2/3 (F) and Smad7 (G) normalized with GAPDH were quantified by densitometry. Data are presented as mean ± SEM for the groups of three animals (1-way ANOVA, followed by Sidak’s post hoc tests). *p < 0.05 vs. LS; **p < 0.05 vs. HS; &p < 0.05 vs. HS+CANA; &&p < 0.05 vs. HS+IRB.

## Discussion

The beneficial renal effects of SGLT2 inhibitors have been observed in patients with nondiabetic nephropathy [11]. However, distinct from clinical trials, it is surprising that results from preclinical studies in rodents with nondiabetic renal injury have thus far been provided rather inconsistently, which is not due to the different SGLT2 inhibitors used. ARBs are widely used to treat hypertension and hypertension-related heart and kidney damage by alleviating tissue fibrosis [12] and improving endothelial function [13]. The results of this study demonstrate that SGLT2 inhibitors and ARBs have nephroprotective effects in the case of SSHT, and greater benefits may be obtained from the combination of the two.

In this study, canagliflozin, irbesartan, and the combined therapy induced various effects on BW, MFI, MWI, SBP, and HR in Dahl SS rats fed an HS diet. An HS diet lowered the BW of Dahl SS rats, and canagliflozin aggravated the weight loss caused by an HS diet, but irbesartan did not. The reduction in BW by SGLT2 inhibitors was consistent with the findings of a previous clinical study [14], which appeared to be related to the SGLT2 inhibitor-decreased glucose level, thus enhancing fat utilization [15]. The effects of SGLT2 inhibitors on MFI are inconsistent due partly to different disease models [16]. Canagliflozin significantly decreased MFI during the present study, while irbesartan did not significantly affect MFI. Our data, together with previously published findings [17], clearly show that an HS diet significantly increased MWI in Dahl SS rats, and canagliflozin can further improve the level of MWI in Dahl SS rats fed an HS diet. However, irbesartan did not affect the HS–induced increase in water intake, indicating that the change in water intake is independent of AT1-receptor stimulation [18]. The urine volume of Dahl SS rats fed an HS diet was significantly increased by canagliflozin, the level of which was alleviated by irbesartan. These results may be explained by the diuretic action of canagliflozin being enhanced by angiotensin II [17].

However, the effects of SLGT2 inhibitors on HS-induced hypertension in rats remain controversial [16, 19]. In the present study, our data showed that there was a nonsignificant trend for SBP lowering by canagliflozin, despite its very pronounced diuresis activity. Irbesartan significantly decreased the SBP of HS Dahl SS rats without increasing urine volume. The SBP of the combination therapy was reduced the most. The nonsignificant decrease in SBP by canagliflozin may partly result from a compensatory activation of sympathetic tone and renin release by an HS diet and natriuresis [19]. Furthermore, although the noninvasive tail-cuff methodology is well accepted for blood pressure monitoring in rats, this method also has some uncertainties due to the need for habituation to restrain and other influences on tail circulation [20].

Contrary to previous studies [21], our data show that canagliflozin treatment significantly increased the blood glucose of Dahl SS rats after a 12-week HS diet, which may be explained by the reduction of blood volume through natriuresis. Development of kidney function in HS Dahl SS rats, including albuminuria, Scr, and FENa, was prevented by irbesartan and the combined treatment but not by canagliflozin. The effects of SGLT2 inhibitors on renal function in our experimental cohorts were consistent with previous clinical [22] and animal [23] studies. Similar to an experiment in Dahl SS diabetic rats [23], maximal renoprotection from kidney function was achieved when canagliflozin was combined with irbesartan, of which the underlying mechanism warrants further consideration. However, inconsistent with previous studies in nondiabetic rats [24], the prevention of BUN and other renal injury markers (Cys-C and KIM-1) in the present study was not observed in different treatment groups.

There have been few reports on the renoprotection of SGLT2 inhibitors in SSHT, and the molecular mechanism of their antifibrotic effects remains unknown. In our experimental conditions, reduction of renal fibrosis determined by Masson staining, when irbesartan was administered alone or in combination with canagliflozin, was observed in line with the improved functional and histological parameters. Moreover, canagliflozin administration prevented the development of renal fibrosis without blood pressure or renal function changes [19]. Indeed, maximal renoprotection from morphologic changes and renal fibrosis was achieved from the combined therapy, which was consistent with a previous finding in Dahl SS diabetic rats [25]. Of further interest, we explored the effect of canagliflozin and irbesartan on TGF-β1/Smad2/3 signaling, which plays a key role in the development of renal fibrosis.

Previous studies have revealed that SGLT2 inhibitors exert a clear protective effect on fibrosis in different tissues through inhibition of the TGF-β/Smad signaling pathway both in vivo and in vitro [26]. Moreover, the expression of TGF-β1 was also suppressed by ARBs to alleviate renal pathological lesions and protect renal function partly independent of AT1R expression [27]. In the present study, TGF-β1, as determined by IF and WB, was also more highly expressed in the kidneys of the HS group than in the kidneys of the LS group, which was accompanied by upregulated levels of fibronectin, collagen I, α-SMA, Smad2/3, and p-Smad2/3. However, both canagliflozin and irbesartan reduced TGF-β1 and other profibrotic factors with a positive correlation with renal fibrosis. Furthermore, we found that the combined therapy had an additive effect on the suppression of profibrotic factors and TGF-β1/Smad2/3 signaling, which was more susceptible to canagliflozin than irbesartan in the kidney. These results suggest that canagliflozin may act directly to decrease renal fibrosis and TGF-β1/Smad2/3 activation in salt-sensitive hypertensive kidneys partially independent of its antihypertensive effects. Taken together, we speculate that an SGLT2 inhibitor may exert a better antifibrotic effect on renal fibrosis in SSHT, which might be related to the suppression of TGF-β1/Smad2/3 signaling.

To our knowledge, this is the first study to compare the difference in renoprotection between canagliflozin and irbesartan in experimental SSHT, including monotherapy and combined therapy. These findings provide novel data supporting that these two medications may exert different effects on physiological indexes, kidney function, renal fibrosis, and the TGF-β/Smad2/3 pathway. The data indicate that irbesartan may attenuate hypertensive renal injury by reducing blood pressure, while canagliflozin exerts a better antifibrotic effect, possibly through inhibition of the TGF-β/Smad2/3 pathway. However, in almost all aspects of renoprotection, the combined therapy was consistently superior to the monotherapy.

However, there are still several limitations in this study. First, the current research is in vivo, and the results were not verified in vitro. Second, the investigation of detailed renal function is insufficient. Third, the mechanism underlying the inhibitory effect on hypertensive renal injury was not fully elucidated. Nevertheless, our research can be considered a starting point for further studies aiming to investigate the difference in renoprotection between SGLT2 inhibitors and ARBs in SSHT.

## Conclusions

In conclusion, the current results indicate that canagliflozin and irbesartan exert different benefits on nephroprotection in SSHT models. Irbesartan may have a better antihypertensive effect, while the antifibrotic effect of canagliflozin may be more remarkable, possibly through inhibition of the TGF-β/Smad2/3 pathway. Consistently, additional molecular and histological benefits were offered when canagliflozin and irbesartan were coadministered, but the nephroprotective cellular mechanisms of the combined therapy remain to be further explored. Taken together, our study suggests that selective SGLT2 inhibitors can be used in combination with ARBs to further prevent the progression of nondiabetic nephropathy, especially in SSHT.

## Materials and methods

### Experimental protocol

All animal experiments in this research were carried out in compliance with the Guide for the Care and Use of Animal Ethics Committee of Hebei Medical University and approved by the ethics committee of Hebei General Hospital (No. 2022086). Fifty-seven seven-week-old male Dahl SS rats were purchased from Beijing Vital River Laboratory Animal Technology Company (Beijing, China). After the pretreatment period at the animal facility of the Clinical Research Center, Hebei General Hospital, all rats were randomly divided into the following five groups according to their diets and treatments: LS group (low salt control group, n=7), LS diet with vehicle treatment (5% hydroxyethylcellulose, 30 mg/kg/d); HS group (HS control group, n=10), HS diet (8% NaCl) with vehicle treatment; HS + CANA group (HS canagliflozin group, n=10), HS diet (8% NaCl) with canagliflozin treatment (canagliflozin, 30 mg/kg/d); HS + IRB group (HS irbesartan group, n=10), HS diet (8% NaCl) with irbesartan treatment (irbesartan, 30 mg/kg/d); HS + CANA + IRB group (HS canagliflozin + irbesartan group, n=10), HS diet (8% NaCl) with canagliflozin and irbesartan (canagliflozin and irbesartan, 30 mg/kg/d, respectively). To achieve better oral administration, all drugs were dissolved in 5% hydroxyethylcellulose (MedChemExpress, 10 mg/ml). Food and water intake were measured daily. Body weight (BW) and SBP were measured weekly. Random blood glucose (RBG) and fasting blood glucose (FBG) were performed at two weeks before the end of the experiment. Twenty-four-hour urine was collected at baseline and the end of the experimental procedure. After being fasted for ~4 h, all rats were anesthetized with sodium pentobarbital (50 mg/kg body weight, i.p.). **Whole blood was** obtained by **the abdominal aorta and** kidneys were isolated.

### Plasma and urine collection and analysis

Before the end of the experiment, 3 rats from each group were housed individually in metabolic cages to collect the 24-hour urine under the conditions of feeding and water consumption. Blood samples were collected from the abdominal aorta, and serum was separated by centrifugation at 3000 rpm for 10 min at 4 °C and stored at − 80 °C. The biochemical indexes in both urine and plasma were determined using an automatic biochemical analyzer.

### Blood pressure measurement

Systolic blood pressure (SBP) was measured by tail-cuff plethysmography (BP-2000; Visitech Systems, Inc.) according to the manufacturer’s instructions. Measurements were obtained for each conscious rat every two weeks until sacrifice. Before each measurement, the rats were prewarmed at 36 °C for 15–20 minutes to obtain reliable results. The average of three pressure readings was calculated for each measurement. SBP was always measured at the same time of day (10-12 am) by the same experimenter.

### Histologic evaluation

One-half of the left kidney from each rat was fixed in 4% paraformaldehyde for 24 hours. After dehydration, they were embedded in paraffin, and 5-μm-thick sections were stained with hematoxylin and eosin (H&E) for morphological analysis and with Masson’s trichrome for evaluation of fibrosis. All sections were analyzed by light microscopy (Nikon, Japan). The renal fibrosis area fractions were quantified using ImageJ software in 10 randomly chosen, non-overlapping fields (400× magnification) for each section.

Immunofluorescence (IF) staining was performed to detect the expression of renal TGF-β1 in rats. After being sealed, the kidney tissue sections were incubated with anti-TGF-β1 antibody (Abcam, USA) overnight at 4 °C at the indicated dilutions. After returning to room temperature, the samples were washed with phosphate-buffered saline (PBS) and incubated with fluorescein-labeled secondary antibody (Zen BioScience, China) for 1 h at 37 °C. Finally, the sections were counterstained with DAPI for nuclear staining and visualized by a fluorescence microscope (Zeiss, Germany). The fluorescence intensity of the positive area was measured by ImageJ software. For each experimental group, five random fields were selected to calculate the mean fluorescence activity.

### Western blot

The protein concentrations of renal samples were determined with a bicinchoninic acid (BCA) protein assay kit (Thermo Fisher Scientific, USA). Protein samples (50 μg) were separated by 10% sodium dodecyl sulfate-polyacrylamide gel electrophoresis (SDS-PAGE) and transferred onto a polyvinylidene difluoride (PVDF) membrane. The membranes were blocked with 5% skimmed milk for 2 h at room temperature and then incubated with primary antibodies overnight at 4 °C as follows: anti-collagen I (ab270993; Abcam, USA; 1:1000), anti-α-smooth muscle actin (α-SMA) (ab7817; Abcam, USA; 1:3000), TGF-β1 (ab179695; Abcam, USA; 1:1000), anti-Smad2/3 (ab202445; Abcam, USA; 1:1000), anti-pSmad2/3 (ab254407; Abcam, USA; 1:1000), anti-Smad7 (sc-265846; Santa Cruz Biotechnology, USA; 1:1000), and anti-glyceraldehyde-3-phosphate dehydrogenase (GAPDH) (ab181603; Abcam, USA; 1:10000 dilution). The membranes were subsequently washed and then incubated with an appropriate horseradish peroxidase (HRP)-conjugated secondary antibody diluted 1:5000 for 1 hour at room temperature. After exposure to hyper film enhanced chemiluminescence (ECL) (Thermo Fisher Scientific, USA), the antigen-antibody complexes were detected using Odyssey XF (LI-COR, USA). The signal intensities of target bands were analyzed using ImageJ software (National Institutes of Health, USA). The optical density of the corresponding internal reference GAPDH was used for correction.

### Statistical analysis

Statistical analysis was performed using IBM SPSS 23.0 software. All data are expressed as mean ± standard error of the mean (SEM). The statistical significance of differences between the groups was determined using one- or two-way analysis of variance (ANOVA) combined with the Newman–Keuls post hoc test. For all tests, a value of P < 0.05 was considered statistically significant.

## Data availability statement

The original data presented in the study are included in the article, and further inquiries can be directed to the corresponding author.

## Funding

This work was supported by a grant from the 2019 Hebei Science and Technology Project (No. 19277787D) and the 2019 Hebei Innovation Capability Promotion Project (No. 199776249D).

## Conflicts of Interest

The authors indicated no potential conflicts of interest.

## Notes

### Competing Interest Statement

The authors have declared no competing interest.

